# CRISPR-BEST: a highly efficient DSB-free base editor for filamentous actinomycetes

**DOI:** 10.1101/582403

**Authors:** Yaojun Tong, Helene L. Robertsen, Kai Blin, Andreas K. Klitgaard, Tilmann Weber, Sang Yup Lee

## Abstract

Filamentous actinomycetes serve as major producers of various natural products including antimicrobial compounds. Although CRISPR-Cas9 systems have been developed for more robust genetic manipulations, concerns of genome instability caused by the DNA double-strand breaks (DSB) and the toxicity of Cas9 remain. To overcome these limitations, here we report development of the DSB-free, single-nucleotide resolution genome editing system **CRISPR-BEST** (**CRISPR**-**<u>B</u>**ase **<u>E</u>**diting **<u>S</u>**ys**<u>T</u>**em). Specifically targeted by an sgRNA, the cytidine deaminase component of CRISPR-BEST efficiently converts C:G to T:A within a window of approximately seven-nucleotides. The system was validated and successfully used in different *Streptomyces* species.

## Main

More than 70% of current antibiotics are derived from natural products of actinomycetes. Genome mining indicates that these organisms still possess a huge unexploited potential of producing our future antimicrobial drugs^1^. However, for exploiting this potential, modern bio-technologies, such as metabolic engineering and synthetic biology, are heavily relying on efficient genetic manipulation or gene editing approaches^1^. Unfortunately, it is relatively difficult to do genome manipulation of actinomycetes, mainly due to their mycelial growth, intrinsic genetic instability and very high GC-content (>70%) of their genomes. There are established traditional mutagenesis methods, but they are relatively inefficient and very time- and labor-consuming^2^.

Recently, more efficient CRISPR-Cas9 systems were developed for scar-less gene knockout, knockin and reversible gene knockdown in actinomycetes^3^. Although these systems provide excellent flexibility and high efficiency, severe challenges still remain. In many actinomycetes, the (over)expression of Cas9 has severe toxic effects and leads to a high number of unwanted off-target effects^3^. Furthermore, the linear chromosomes show a relatively high intrinsic instability and can tolerate large-scale chromosomal deletions and rearrangements^4^. DNA double-strand breaks (DSB) in the arm region are considered major triggers of this instability^5^ and often co-occur with the mutagenesis procedures.

Here, we present an alternative highly efficient approach to generate mutations in filamentous actinomycetes without the requirement of DSBs. The targeted conversion of cytidine (C) to thymidine (T) can lead to the introduction of stop codons^6-9^ and loss-of-function mutations into the coding genes of different organisms. In particular, we also can introduce rare TTA codons to artificially put genes under BldA control^10^. Such tools are called “base editors”. A prominent example is the BE3 system for editing human cell lines^11^, which was constructed by artificially fusing the rat APOBEC1 (rAPOBEC1) cytidine deaminase, a Cas9 nickase (Cas9n) and a uracil glycosylase inhibitor (UGI). Localized by the target binding capability of sgRNA/Cas9n, the deamination reaction takes place in the single strand DNA within R-loop of the sgRNA:target DNA complex. The deamination of the targeted C in a C:G base pair results in a U:G mismatch (Fig. 1a, and 1c). As U is an illegitimate DNA base, it normally will be recognized and then excised by uracil-DNA glycosylases (UDGs)^12^. This initiates the conserved nucleotide excision repair (NER)^13^, leading to the reversion to the original C:G base pair. However, this process can be inhibited by UGI. This triggers the conserved cellular mismatch repair (MMR)^14^ to efficiently convert the U:G to a U:A base pair. The efficiency of the MMR repair can be increased by introducing a single-stand DNA nick in proximity to the editing site^11^. Thus, when using BE3 or related systems, the resulting G:U mismatched base pair is retained and converted into an A:T base pair upon Cas9n-mediated nicking of the G-containing DNA strand followed by DNA synthesis. This process generates permanent modifications of the target DNA without the requirement of DSB. By clever selection of the target sites, base editors can thus either generate point mutations resulting in amino acid replacements or the introduction of STOP codons (Fig. 1e, and Supplementary Table 6).

**Fig. 1.**
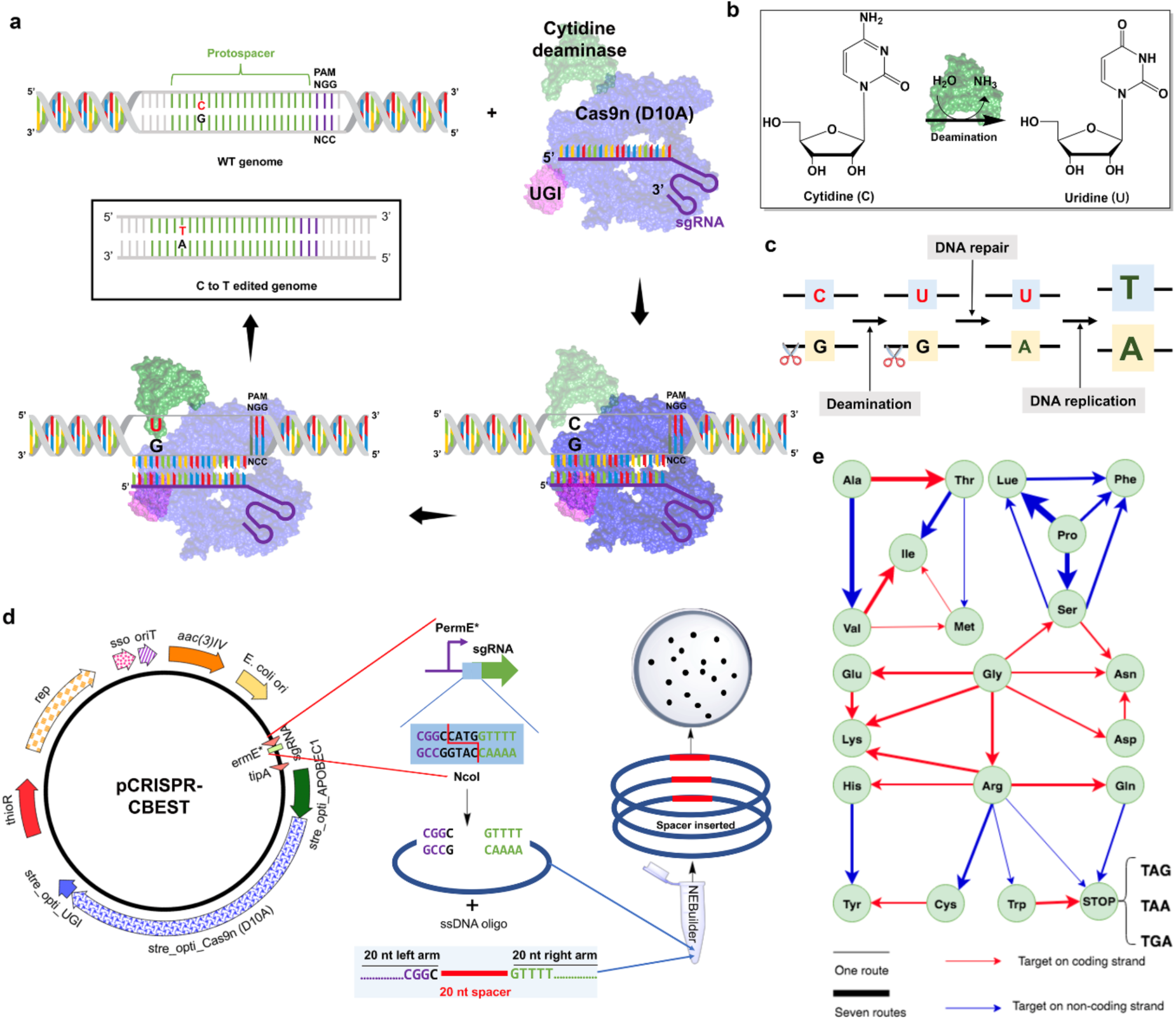
Rationale and workflow CRISPR-BEST. **a.** Overview of the base editing strategy. The target C within the editing window is indicated in red, the possible active domains in each step is shown in a brighter color. Firstly, sgRNA (purple) binds to D10A Cas9n (blue), ending up with Cas9n:sgRNA complex. Secondly, the Cas9n:sgRNA complex finds and binds its target DNA, which mediates the separation of the double-stranded DNA to form the R-loop structure. Thirdly, a tethered *Streptomyces* optimized cytidine deaminase rAPOBEC1 (green) converts the target C in the non-targeted strand to a U by cytidine deamination. Lastly, the resulting U:G heteroduplex is permanently converted to a T:A base pair. **b.** The enzymatic reaction of the cytidine deamination process. **c.** Detailed conversion process of a C:G base pair to a T:A base pair. Due to the inhibition of the nucleotide excision repair (NER) pathway by UGI, the cellular mismatch repair (MMR) becomes the dominant DNA repair pathway. It preferentially repairs the mismatch in a nicked strand. Therefore, the G in the targeted strand, which is nicked by D10A Cas9n, is going to be efficiently replaced by A and in the next replication cyclerepaired to a T:A base pair. **d.** The CRISPR-BEST plasmid is a pSG5 replicon based, temperature sensitive, *E. coli*-*Streptomyces* shuttle plasmid. *S. coelicolor* A3(2) codon optimized rAPOBEC1, Cas9n (D10A), and UGI were fused together, and can be expressed under control of the leaky *tipA* promoter. The sgRNA cassette is under control of the *ermE*\* promoter. A PCR-free, one-step ssDNA bridging approach can be applied for the 20bp-spacer cloning. **e.** Representation of the possible amino acid exchanges resulted by CRISPR-BEST. Blue lines indicate that the target C is in coding strand, while red lines indicate that the target C is in non-coding strand. The thickness of the lines indicates the number of possible routes that can end up with the same amino acid exchange by CRISPR-BEST.

In order to address the limitations of CRISPR-Cas9 in filamentous actinomycetes, here we report development of a DSB-free, single base pair editing system termed as **CRISPR-BEST**: **CRISPR**-**<u>B</u>**ase **<u>E</u>**diting **<u>S</u>**ys**<u>T</u>**em. As core components, the gene encoding the cytidine deaminase rAPOBEC1 (genbank: NM_012907.2) was codon optimized for *Streptomyces* and fused to the N-terminus of Cas9n (D10A) using a 16-amino acid flexible linker. In order to inhibit the NER, a *Streptomyces* codon optimized UGI (genbank accession number: YP_009283008) was fused to the C-terminus of Cas9n by a short linker. sgRNAs can be introduced using a highly effective single strand DNA oligo bridging method. For details, please see Online Methods.

For a proof-of-concept, the actinorhodin biosynthetic gene cluster region of *S. coelicolor* A3(2) was selected as a target. Potential protospacers containing the editable cytidines were identified in the genes encoded in the target region using the updated CRISPy-web (https://crispy.secondarymetabolites.org), the updates now make the sgRNA identification tool CRISPy-web^15^ directly support CRISPR-BEST sgRNA design. In total, twelve protospacers were selected to construct sgRNAs, six targeting the coding strand and six targeting the non-coding strand.

The reported base editors have a less than ten-nucleotide editing window^11, 16^. Therefore, we investigated all the cytidines within the 10-nucleotides in the PAM-distal region. We observed that not a single cytidine was converted into a thymidine in the first three nucleotides of the hypothetic editing window of all twelve protospacers (Supplementary Fig. 1). Thus, the editing window of CRISPR-BEST can be assigned to seven nucleotides (positions 4 to 10 in the hypothetic editing window) in the PAM-distal region (Fig. 2b). The cytidines in the editing window were converted into thymidines with frequencies between 30% and 100% (Supplementary Fig. 1). Only in three cases, where the C is preceded by a G, no conversion was observed.

**Fig. 2.**
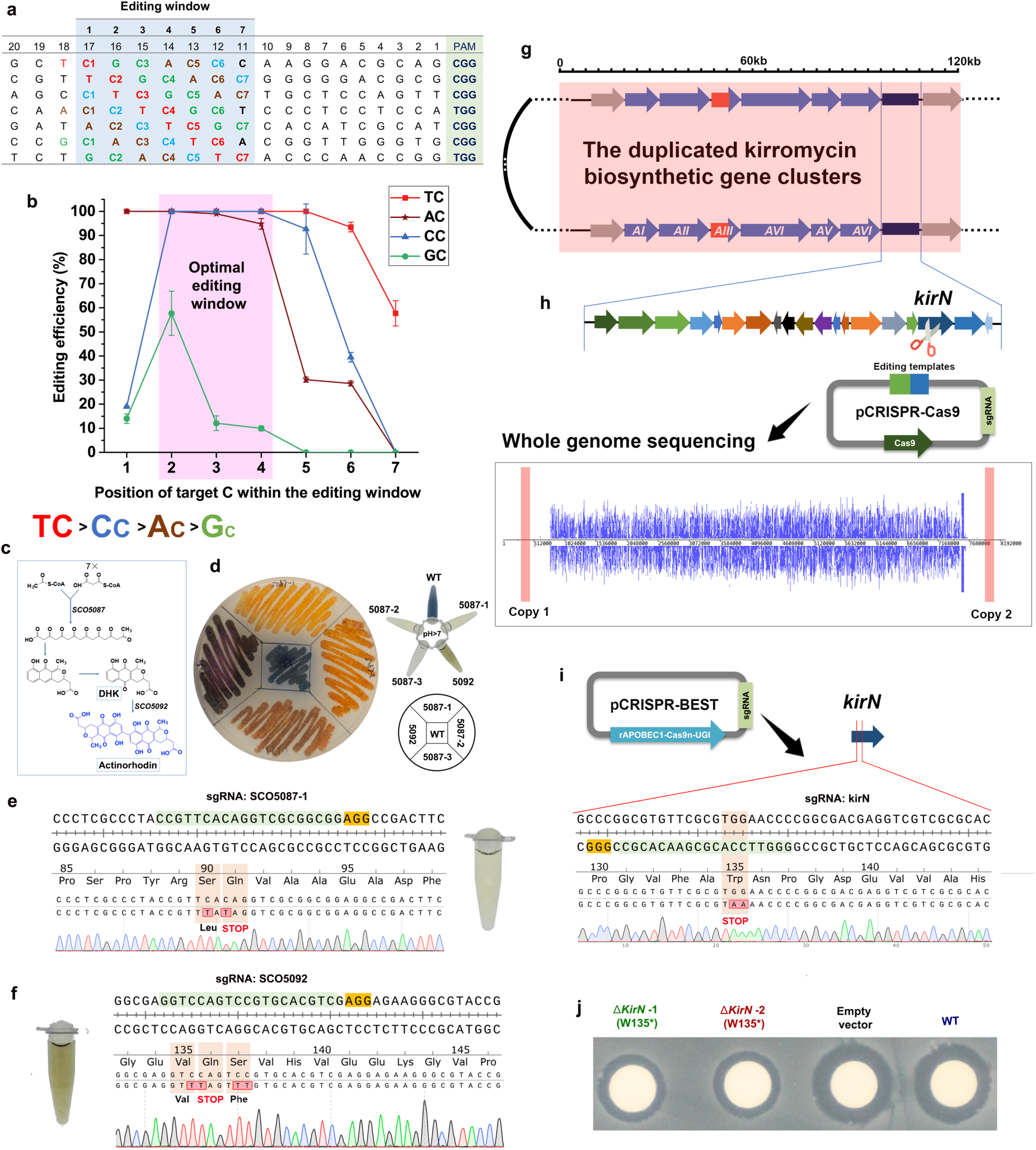
CRISPR-BEST characterization and applications. **a.** Positional effect of each NC combination on editing efficiency *in vivo.* A matrix of TCGCACC was designed to investigate the optimal NC combination and target C position within the editing window. 20nt protospacer and its PAM was displayed. The editing window was masked in light blue. **b.**Each NC combination was varied from positions one to seven within the protospacer. The target regions of 10 to 20 CRISPR-BEST treated exconjugants of each protospacer were PCR amplified and Sanger sequenced. For mixed trace signals, the secondary peak calling function of CLC Main Workbench 8 (QIAGEN Bioinformatics, Germany) was applied to calculate the editing efficiency. The 3-nt window in pink showed the optimal editing efficiency. Values and error bars were the mean and standard deviation of two to three independent conjugations using the same pCRISPR-BEST plasmids. **c.** A simplified biosynthetic route of the blue-pigmented polyketide antibiotic actinorhodin. SCO5087, coding for the actinorhodin polyketide beta-ketoacyl synthase subunit alpha, and SCO5092, coding for the actinorhodin polyketide dimerase were selected as editing targets. **d.** One *S. coelicolor* A3(2) WT, three base-edited SCO5087 mutants (ΔSCO5087 (Q91*), SCO5087 (R89C, S90L), and SCO5087 (R89C)), one base-edited SCO5092 mutant (ΔSCO5092 (Q136*)) were streaked onto ISP2 agar plate with apramycin. The same corresponding extracts were shown as well. **e.** and **f.** Sanger sequencing traces of the region containing a protospacer together with its PAM. Protospacers are highlighted in light green, PAM sequences in yellow, the codons and corresponding amino acids are indicated, detailed editing efficiency were shown in Supplementary Fig. 2a. **g.** Schematic representation of the linear chromosome of *S. collinus* Tü365, in which two copies of kirromycin biosynthetic gene cluster located approximately 341 kb from the left, while 422 kb from the right end of the chromosome are shown. Kirromycin is a narrow-spectrum antibiotic. Its biosynthesis is encoded by two identical 82 kb gene clusters that are located in the long inverted repeats of the chromosome arms^19, 20, 24^. Within the kirromycin BGC, *kirN* codes for an enzyme that is very similar to primary metabolism CCR crotonyl-CoA reductase/carboxylases (CCR)^20, 24^ and thus it is speculated that it is involved in enhancing the pool of ethylmalonyl-CoA, one building block of kirromycin.A key module containing the *kirN* gene was zoomed in as indicated. **h.** CRISPR-Cas9 based homologous recombination approach was unsuccessfully used to generate an in-frame Δ*kirN* mutant. Paired-stack view of Illumina MiSeq reads for the CRISPR-Cas9 Δ*kirN*-mutant mapped against the reference genome of *S. collinus* Tü365. Mapping results showed that both kirromycin cluster encoded near the chromosome ends were lost. The deletion comprises 787,795 bp from the left and 630,478 bp from the right end. **i.** CRISPR-BEST was used to generate *kirN* null mutant by a STOP codon introduction. Validation of the correct editing of *kirN*_*W135→STOP*_ by Sanger sequencing of PCR amplified target region. **j.** Bioactivity testing of four extracts from WT, empty vector (no spacer), and two clones of CRISPR-BEST edited *kirN*_*W135→STOP*_ using *Bacillus subtilis* 168 as indicator strain.

In previous studies it was demonstrated that the rAPOBEC1 based base editor showed different performance when used *in vitro* compared with *in vivo*^11^. In order to systematically evaluate the effects of sequence context and the target C position on editing efficiency in a “close to application” context *in vivo*, we designed a matrix based on the four possible combinations of C with the other three nucleotides, A, T, and G (TCGCACC). In the matrix, the target C of each NC combination was distributed in all seven possible positions (Fig. 2a). We used PatScanUI^17^ to identify the possible protospacer variants in the genome of *S. coelicolor* A3(2). Seven protospacers in nonessential genes were selected and tested experimentally (Fig. 2a). By calculating the C→T conversion efficiency (Fig. 2b), it became obvious that the CRISPR-BEST system is accepting its deamination substrates in the priority of TC>CC>AC>GC (Fig. 2b). This finding is consistent with other reports^11, 18^. Within the seven-nucleotide editing window, we observed that positions 2, 3 and 4 showed highest editing efficiency (Fig. 2b).

By converting C to T in any of the 64 natural codons, 32 different amino acid substitutions can be generated, which cover almost all 20 natural amino acids (Fig. 1e). As an application, Arg codons (**C**GA), Gln codons (**C**AA and **C**AG), and Trp codons (T**GG**, target C in non-coding strand) are particularly interesting, as they can be converted to stop codons (TGA, TAA, and TAG) by cytidine deamination. For generalizing this strategy, we systematically analyzed the number of potential target sites that lead to STOP codon introduction into the nonessential secondary metabolites biosynthesis genes of the model actinomycete *S. coelicolor* A3(2) and non-model actinomycete *Streptomyces collinus* Tü365 using the updated CRISPy-web. An average of about 13 and 14 possible target sites per gene were identified for *S. coelicolor* A3(2) and *S. collinus* Tü365, respectively (Supplementary Table 7 and 8). To validate CRISPR-BEST on amino acid substitutions *in vivo*, two genes, SCO5087 (ActIORF1, KSα of minimal PKS) and SCO5092 (ActVB, dimerase), from the biosynthetic pathway of the diffusible, blue-pigmented polyketide antibiotic actinorhodin in *S. coelicolor* (Fig. 2c) were selected. sgRNAs targeting these two genes were designed and cloned into CRISPR-BEST plasmids. Sanger sequencing of the targeted region revealed that all target cytidines were converted to thymidines, ending up with desired amino acid substitutions (Fig. 2d-2f, Supplementary Fig. 2a and 2b) or introduction of STOP codons (Fig. 2e and 2f). The loss-of-function of the gene encoding the actinorhodin polyketide beta-ketoacyl synthase subunit alpha ActIORF1 (SCO5087) completely eliminates actinorhodin biosynthesis (Fig. 2c) and thus the dark-blue colored phenotype of the colonies (Fig. 2d). While ActVB (SCO5092) catalyzes the dimerization of two polyketide precursors as one of the last steps of the actinorhodin biosynthesis, a null mutant (Fig. 2f) in this gene leads to the accumulation of the intermediate dihydrokalifungin (DHK) (Fig. 2c). Compared to actinorhodin, the colonies exhibit brownish color on ISP2 agar plate (Fig. 2d). In all four tested cases, the targeted C were converted to T with an editing efficiency of nearly 100% (Supplementary Fig. 3).

To include a “real world” test example, we next elucidated if CRISPR-BEST is capable of simultaneously inactivating two identical gene copies of the gene *kirN* (Locus B446_01590 and B446_33700) in the duplicated kirromycin biosynthetic gene clusters (BGCs)^19^ of the non-model actinomycete strain *S. collinus* Tü365 (Fig. 2g). When using the classical CRISPR/Cas9 system, all clones obtained after pCRISPR-Cas9 treatment targeting *kirN* completely lost kirromycin production (Supplementary Fig. 4b). Further investigation revealed that the complete loss of kirromycin production and unsuccessful complementation with plasmid-encoded *kirN* was due to large deletions of both chromosome arms (787,795 bp from the left arm, 630,478 bp from the right arm), which contain the two copies of the kirromycin BGC (Fig. 2h). These deletions were likely caused by the simultaneous DSBs introduced by Cas9.

For CRISPR-BEST, a protospacer within *kirN* was identified that should introduce an early STOP codon (Fig. 2i). After transferring the CRISPR-BEST plasmid with the *kirN*-targeting sgRNA into *S. collinus* Tü365, sequencing of PCR products of the target region demonstrated that the cytidines were converted to thymidines and thus a STOP codon was successfully incorporated into *kirN* (Fig. 2j). In production assays using the *kirN*_*W135→STOP*_ mutant, kirromycin was still produced but with a much lower yield compared to the wild-type strain (WT) (Supplementary Fig. 4a, 4c-4e), which is consistent with our previous observation of the mutant that was generated by classic homologous recombination-based gene knockout approach^20^.

The above examples clearly demonstrate the potential of CRISPR-BEST in efficiently introducing mutations in the actinomycetes genome without involving DSBs, resulting in reduced risk of genome instability often caused by CRISPR-Cas9. A comprehensive comparison of CRISPR-BEST with CRISPR-(d)Cas9 or classical actinomycete mutagenesis approaches is included in Supplementary Table 1. Taken together, CRISPR-BEST is a powerful addition to the actinomycete CRISPR-Cas9-based genome editing toolbox.

## Methods

### Strains, plasmids, and culture conditions

The strains and plasmids used in this study are listed in Supplementary Table 2. All plasmids were maintained in *E. coli* DH5alpha. All *E. coli* strains and *Bacillus subtilis* 168 were grown in LB medium (agar and liquid) at 37°C. *Streptomyces* strains were grown at 30 °C in either ISP2 (Yeast extract 4 g/l, Malt extract 10 g/l, Dextrose 4 g/l, 20 g/l Agar is added for solidification) for seed culture and DNA preparation, or in MS-MgCl_2_ (20 g/l each D-mannitol, soya flour, agar, and 10mM MgCl_2_) for sporulation and conjugation. Kirromycin production medium (10 g/l full-fat soy flour, 10 g/l D-mannitol, and 5 g/l CaCO_3_, dissolved in tap water and pH adjusted to 7.4 prior to autoclaving) was used for kirromycin production assays. Appropriate antibiotics were supplemented as necessary (50 μg/ml apramycin; 50 μg/ml nalidixic acid; 0.5 μg/ml thiostrepton; 25 μg/ml kanamycin; and 25 μg/ml chloramphenicol). *E. coli* ET12567/pUZ8002 was used for conjugating plasmids into streptomycetes as described previously^2^.

### Construction of CRISPR-BEST plasmids

All primers used in this study are listed in Supplementary Table 3.

A self-replicating pSG5-based thermosensitive *E.coli-Streptomyces* shuttle vector pGM1190^21^ (Fig. 1b) was used as the backbone plasmid to construct the CRISPR-BEST plasmid. The sgRNA cassette design is similar to our previous pCRISPR-Cas9 system^22^. In order to simplify the 20nt-spacer cloning process and increase its cloning efficiency, we modified the original sgRNA cassette to be compatible with single strand DNA (ssDNA) oligo bridging method (lower-middle panel of Fig. 1d) by removal of a G from the pGM1190-sgRNA^22^ plasmid with primers removalG_F and removalG_R, resulting in plasmid pGM1190-sgRNAnoG. The transcription of the sgRNA is controlled by a constitutive promoter *ermE\**, and terminated by a *to* terminator. Due to the huge differences of codon usage between streptomycetes and other organisms, the cytidine deaminase rAPOBEC1 (apolipoprotein B mRNA editing enzyme catalytic subunit 1 from *Rattus norvegicus*, genbank accession number: NM_012907.2), the Cas9n (D10A), and the UGI from *Bacillus* phage AR9 (genbank accession number: YP_009283008) were codon optimized to *S. coelicolor* A3(2) using Genscript’s OptimumGene™ algorithm (Supplementary Fig. 5) and then synthesized by Genscript. The stop codon removed rAPOBEC1 was fused to the N-terminus of the start and stop codons removed Cas9n (D10A) using a 16-amino acid flexible linker (SGSETPGTSESATPES, the encoding DNA sequence was also Streptomyces codon optimized). The start codon removed UGI was then fused to the C-terminus of Cas9n (D10A) by a SGGS linker. Gibson assembly was used to assemble the DNA fragment encoding the N-rAPOBEC1-linker-Cas9n-linker-UGI-C fusion protein into *Nde*I and *Xba*I digested pGM1190-sgRNAnoG plasmid, the fusion protein is under control of the thiostrepton inducible *tipA* promoter, resulting in the final pCRISPR-BEST plasmid, which is been depositing to Addgene.

### Single-strand DNA based PCR-free spacer cloning protocol

To use the pCRISPR-BEST for base editing applications, only one step is required, which is the insertion of a 20nt spacer into the sgRNA scaffold. A ssDNA oligo based, PCR-free method was adopted for spacer cloning in this study. The oligo was designed as CGGTTGGTAGGATCGACGGC<u>**N20**</u>GTTTTAGAGCTAGAAATAGA. As designed, the pCRISPR-BEST plasmid can be linearized by *Nco*I. By mixing the linearized pCRISPR-BEST plasmid and chemically synthesized spacer containing oligo with the NEBuilder (New England Biolabs, USA). The linearized pCRISPR-BEST plasmid then will be bridged by the spacer containing oligo, ending up with the desired pCRISPR-BEST. Mach1™-T1^R^ *E. coli* (Life Technologies, UK) was used for cloning. Because of the high bridging efficiency, 4-8 clones were directly sanger sequenced using primer “stre_spacer_seq” to screen for the correct constructs. All plasmids generated and used were listed in Supplementary Table 2.

### *In vivo* spacer-matrix design using PatScan

As two key components of this spacer-matrix are the positions and the variants of TCGCACC in the 23nt protospacer plus PAM sequence. The pattern of the matrix is N_2-3_(TC_n_GC_n_AC_n_C_n_)N_12-11_GG, where n = 1 to 7, therefore, the matrix contains in total seven pieces of protospacer (Fig. 2a and Supplementary Table 4). PatScanUI^17^ (https://patscan.secondarymetabolites.org) was used to locate all possible protospacers in the genome of *S. coelicolor* A3(2). Each found protospacer was cross-compared with all spacers of *S. coelicolor* found by CRISPy-web^15^, then the ones with less off-target effects were manually checked if they are inside of essential genes or not, based on the genome annotation of *S. coelicolor* A3(2). The rules for matrix protospacer selection are: not in essential gene; not located too close to the chromosome end; less off-target effects; and if possible, select the ones sharing the same PAM sequence.

### CRISPR-BEST support in CRISPy-web

For the updated CRISPy-web, sgRNAs are identified using the regular CRISPy-web algorithm published previously^15^. All sgRNAs in the region of interest where the potential edit window overlaps with an annotated CDS region are then selected for CRISPR-BEST analysis. The CDSs with overlap to the sgRNA edit windows are split into individual codons. The codons are filtered for overlaps with the edit window again. For sgRNAs on the same strand as the CDS, all possible C to T mutations are recorded, for sgRNAs on the opposite strand, all possible G to A mutations are recorded. Non-conservative mutations changing the encoded amino acid are finally reported in the CRISPy-web interface.

### CRISPR-BEST compatible protospacers identification using CRISPy-web

The procedure is based on our previous report^15^. Briefly, a custom genome or an antiSMASH generated job id needs to be uploaded to CRISPy-web (https://crispy.secondarymetabolites.org). Taking *kirN* of *S. collinus* Tü365 as an example (Supplementary Fig. 7a), all possible protospacers from both DNA strands will be displayed for the *kirN* gene (Supplementary Fig. 7b). By choosing the “Show CRISPR-BEST output” box, all CRISPR-BEST compatible protospacers will be displayed (Supplementary Fig. 7c), showing the possible amino acid substitutions. By subsequentially choosing the “Show only STOP mutations” box, all possible STOP codon introductions will be displayed (Supplementary Fig. 7d), and the selected protospacers can be downloaded as CSV file by clicking the shopping basket located in the up-right corner.

### In-frame deletion of *kirN* using CRISPR-Cas9 based homologous recombination strategy

The in-frame deletion of *kirN* in *S. collinus* Tü365 using CRISPR-Cas9 based homologous recombination approach was carried out as we described in^23^. USER cloning approach was used for the plasmid assembly^23^. The 20nt spacer region GATCGCATTTCGCCAACTAC that specifically targeted on *kirN* was predicted CRISPy-web^15^ (https://crispy.secondarymetabolites.org). The 462 bp sgRNA-*kirN* cassette was ordered as a gBlocks® Gene Fragment from IDT (Integrated DNA Technologies, US) and the full sequence can be found in Supplementary Table 5. The directional assembly of the sgRNA and the two 1kb editing templates, interspaced by the *ermE\** promoter, in the linearized pCRISPR-USER-Cas9 was ensured by the uracil-containing overhangs generated by PCR amplification with primer pair pHR1/pHR2 for the sgRNA gBlocks® gene fragment, pHR3/pHR4 for the *ermE*\* promoter, and primer pairs pHR5/pHR6 and pHR7/pHR8 for the 1kb editing templates up- and down-stream of *kirN*, respectively. The 1 kb editing templates were amplified from genomic DNA of *S. collinus* Tü365.

Upon the USER assembly, correct clones of pCRISPR-Δ*kirN* were identified with control PCR with pHR9/pHR10 and confirmed by Sanger sequencing with primers pHR9 and pHR13. The resulting pCRISPR-Δ*kirN* was introduced into *S. collinus* Tü365 by intergeneric conjugation following a protocol reported previously^2^.

### Base pair changes were evaluated by Sanger sequencing

First, primers that can amplify a several-hundred base pairs region containing the base editing window were designed (Supplementary Table 3). Secondly, colony PCR approach was used to amplify the designed regions directly from streptomycetes colonies. The protocol was modified from our previous publication^22^: about four-square-millimeter actively growing mycelia (for example, 3-day old *S. coelicolor*, before sporulation) of the selected colonies were scraped from the agar plate using a sterile toothpick into 20 μl pure DMSO in PCR tubes. The tubes were shaken and boiled vigorously for 20 min at 100 °C in a heating block. After cooling down to room temperature, the solution was centrifuged at top speed for 30 seconds, 1 μl of the supernatant was used as the PCR template in a 20 μl-reaction with Q5 High-Fidelity DNA Polymerase (New England Biolabs, US). Lastly, the PCR products were cleaned up by kits from Thermo Fisher Scientific, USA and then sanger sequenced by Mix2Seq kit (Eurofins Genomics, Germany).

### Kirromycin fermentation and analysis

The protocol was modified from^20^. Four-day old seed cultures (grown in ISP2), normalized according to wet weight, were inoculated into kirromycin production medium ending up with in total 50 ml in shake flasks. The fermentations were carried out for six days at 30°C in a rotary shaker at 180 rpm. 30 ml of each culture was extracted with 1:1 ethyl acetate for 2 h at room temperature. The extracts were then dried, re-dissolved in 200 μl methanol, and stored in −20°C for further applications.

LC-MS analysis was performed using an ultra-high-performance liquid chromatography (UHPLC) UV/Vis diode array detector (DAD) high-resolution mass spectrometer (HRMS) Orbitrap Fusion mass spectrometer connected to a Dionex Ultimate 3000 UHPLC pumping system (Thermo Fisher Scientific, USA). UV-Vis detection was done using a DAD-3000 set to the range 190 – 700 nm. Injections of 3 μL of each sample was separated using an Acquity UPLC HSS T3 column (2.1 × 100 mm, 1.8 μm) (Waters, USA) at a flow rate of 0.4 mL/min, and a temperature of 30.0 °C. Mobile phases A and B were 0.1 % formic acid in water and acetonitrile, respectively. Elution was performed with a 30 min multistep system. After 5 % B for 1 min, a linear gradient started from 5 % B to 100 % B in 21 min, which was held for another 5 min and followed by re-equilibration to 5 % B until 30 min. HRMS was performed in separate ESI+ and ESI-experiments with a in the range (*m/z*) 200-2,000 at a resolution of 120,000, RF Lens 60 %, and AGC target 5.0e4.

Data analyses were performed with the software Xcalibur 3.1.2412.17 (Thermo Fisher Scientific, USA).

### Bioactivtivity assay of kirromycin

Wild type *Bacillus subtilis* was used as indicator strain. An overnight *B. subtilis* colony of approximately four-square-millimeter was transferred from LB agar plate into 1 ml LB liquid medium in a 1.5 ml Eppendorf tube. The suspension was mixed by vortexing. 200 μl of the above suspension was plated onto a LB agar plate, air drying for 5 min inside of a clean bench. Sterilized paper disks were placed onto the same LB agar plate, then 20 μl of each exact was added onto the paper disks. The resulting LB plate was incubated at 37°C incubator for 24 h, the image was taken by a ColonyDoc-It™ Imaging Station (Analytik Jena AG, Germany).

### Assay for actinorhodin extraction

Exconjugants were picked and streaked onto apramycin containing ISP2 agar plate and incubated for five days at 30 °C. The photos of related plates were taken by a ColonyDoc-It™ Imaging Station (Analytik Jena AG, Germany). Actinorhodin extraction assay was carried as following procedural: 10 ml of seven days old *S. coelicolor* ISP2 culture was mixed 1:1 with 1 N NaOH. Extracting for 4 h using a magnetic stirrer at room temperature. The suspensions were centrifuged at 10,000 g for 5 min, the supernatants were transferred into PCR tubes. All tubes were placed under the same filed for photo taken by a ColonyDoc-It™ Imaging Station (Analytik Jena AG, Germany), so that their color could be directly compared.

### DNA manipulation

All primers and spacers used in this work are listed in Supplementary Table 3 and Supplementary Table 4, respectively. All kits and enzymes were used according to the manufacturers’ recommendations. Standard protocols were used for DNA purification, PCR, and cloning, unless the modifications were indicated. PCR was performed using Phusion High-Fidelity PCR Kit (Thermo Fisher Scientific, US), and Q5 High-Fidelity DNA Polymerase (New England Biolabs, US). Digestion was carried out using FastDigest restriction enzymes (Thermo Fisher Scientific, US). Cloning was carried out using the Gibson Assembly® Master Mix kit and NEBuilder® HiFi DNA Assembly kit (New England Biolabs, US). Genomic DNA was prepared by the Blood & Cell Culture DNA Kit (QIAGEN, Germany). Mix2Seq kit (Eurofins Genomics, Germany) was used for Sanger sequencing.

### Illumina whole genome sequencing and analysis

Illumina sequencing was carried out as we described before^22^. Briefly, a 10 ml five days old *S. collinus* cluture was used for genomic DNA isolation. The genomic library was generated using the TruSeq ®Nano DNA LT Sample Preparation Kit (Illumina Inc., US). The reads obtained from the Illumina sequencing were mapped to the WT *S. collinus* Tü365 reference genome (NCBI accession: CP006259) using the software BWA with the BWA-mem algorithm. The data was inspected and visualized using readXplorer and Artemis.

## Data availability

Source data for Figs. 1e and Fig. 2b and for Supplementary Fig. 5 is available online. Other data is available in the NCBI under accessions (XXXX) and also from the corresponding author upon request.

## Acknowledgements

This work was supported by grants from the Novo Nordisk Foundation (NNF10CC1016517, NNF15OC0016226, NNF16OC0021746). S.Y.L. was also supported by the Technology Development Program to Solve Climate Changes on Systems Metabolic Engineering for Biorefineries (NRF-2012M1A2A2026556 and NRF-2012M1A2A2026557) from the Ministry of Science and ICT through the National Research Foundation (NRF) of Korea.

## Author Contributions

Y.T. conceived the study and designed the experiments. Y.T., H.L.R., and A.K.K., performed laboratory experiments. Y.T., and A.K.K. performed data analysis. K.B., and T.W. designed the spacer identification software. Y.T., T.W., and S.Y.L. wrote the manuscript.

