## Supplementary Information for "CRISPR-BEST: a highly efficient DSB-free base editor for filamentous actinomycetes"

Supplementary Table 1[a]. A direct comparison between CRISPR-BEST and CRISPR-(d)Cas9

| | CRISPR-Cas9 | CRISPR-dCas9 (CRISPRi) | CRISPR-BEST (Base editor) |
|---|---|---|---|
| **Conjugation efficiency**[b] | Medium | Medium | High |
| **Editing efficiency**[c] | High | High | High |
| **Spacer cloning (day)** | 3-5 | 3-5 | 1-2 |
| **Editing templates cloning (day)** | 8-15 | Not required | Not required |
| **Null mutant generation** | Yes | No | Yes |
| **Stress on chromosome** | High | Low | Low |
| **SNP generation** | Hard | No | Easy |
| **Editing repeats** | No[d] | Yes | Yes |

a: Information of this table compiled from [21] and this study; b: Please refer to Supplementary Figure 6; c: Editing efficiency of CRISPR-(d)Cas9 is sgRNA dependent[21], editing efficiency of CRISPR-BEST is also sgRNA dependent, but additional constraints apply (see main text and Fig. 2a and 2b); d: Only one case was tested in this study. It might be strain-dependent.

Supplementary Table 2. Strains and plasmids used/generated in this study

| Strains | Description | Source |
|---|---|---|
| *E. coli* DH5alpha | For routine plasmids maintenance and cloning | New England Biolabs |
| *E. coli* Mach1™-T1R | For routine plasmids maintenance and cloning | Thermo Fisher Scientific |
| *E. coli* ET12567/pUZ8002 | For conjugating plasmids into streptomycetes | Maintained in lab |
| *Bacillus subtilis* 168 | For kirromycin bioactivity testing | ATCC |
| *S. coelicolor* A3(2) | Wild type strain of *S. coelicolor* | Maintained in lab |
| ΔSCO5087 (Q91*) | Base edited *S. coelicolor* A3(2), a stop codon was introduced in SCO5087 | This study |
| SCO5087 (R89C, S90L) | Base edited *S. coelicolor* A3(2), two amino acids were exchanged in SCO5087 | This study |
| SCO5087 (R89C) | Base edited *S. coelicolor* A3(2), a amino acid was exchanged in SCO5087 | This study |
| ΔSCO5092 (Q136*) | Base edited *S. coelicolor* A3(2), a stop codon was introduced in SCO5092 | This study |
| *S. collinus* Tü365 | Wild type strain of S. collinus | Maintained in lab |
| Δ*kirN*-Cas9 | *S. collinus* Tü365 with pCRISPR-ΔkirN | This study |
| ΔkirN (W135*)-1 | Base edited *S. collinus* Tü365, a stop codon was introduced in KirN, clone no. 1 | This study |
| ΔkirN (W135*)-2 | Base edited *S. collinus* Tü365, a stop codon was introduced in KirN, clone no. 2 | This study |
| Tü365_Empty vector | *S. collinus* Tü365 with pCRISPR-BEST without any spacer coloned | This study |
| **Plasmids** | **Description** | **Source** |
| pJET1.2 | For cloning PCR fragments of Sanger sequencing | Thermo Fisher Scientific |
| pCRISPR-USER-Cas9 | Empty vector, USER cloning compatible pCRISPR-Cas9 | [22] |
| pCRISPR-Δ*kirN* | pCRISPR-Cas9 with KirN spacer and editing templates for KirN deletion | This study |
| pCRISPR-BEST | Empty vector, cytidine deaminase based base editor | This study |
| pCRISPR-BEST-5087-1 | pCRISPR-BEST with sgRNA:SCO5087-1 | This study |

| | | |
|---|---|---|
| pCRISPR-BEST-5087-2 | pCRISPR-BEST with sgRNA:SCO5087-2 | This study |
| pCRISPR-BEST-5087-3 | pCRISPR-BEST with sgRNA:SCO5087-3 | This study |
| pCRISPR-BEST-5092 | pCRISPR-BEST with sgRNA:SCO5092 | This study |
| pCRISPR-BEST-Matrix_1 | pCRISPR-BEST with sgRNA:Matrix1 | This study |
| pCRISPR-BEST-Matrix_2 | pCRISPR-BEST with sgRNA:Matrix2 | This study |
| pCRISPR-BEST-Matrix_3 | pCRISPR-BEST with sgRNA:Matrix3 | This study |
| pCRISPR-BEST-Matrix_4 | pCRISPR-BEST with sgRNA:Matrix4 | This study |
| pCRISPR-BEST-Matrix_5 | pCRISPR-BEST with sgRNA:Matrix5 | This study |
| pCRISPR-BEST-Matrix_6 | pCRISPR-BEST with sgRNA:Matrix6 | This study |
| pCRISPR-BEST-Matrix_7 | pCRISPR-BEST with sgRNA:Matrix7 | This study |
| pCRISPR-BEST-spacer_1 | pCRISPR-BEST with sgRNA:spacer1 | This study |
| pCRISPR-BEST-spacer_2 | pCRISPR-BEST with sgRNA:spacer2 | This study |
| pCRISPR-BEST-spacer_3 | pCRISPR-BEST with sgRNA:spacer3 | This study |
| pCRISPR-BEST-spacer_4 | pCRISPR-BEST with sgRNA:spacer4 | This study |
| pCRISPR-BEST-spacer_5 | pCRISPR-BEST with sgRNA:spacer5 | This study |
| pCRISPR-BEST-spacer_6 | pCRISPR-BEST with sgRNA:spacer6 | This study |
| pCRISPR-BEST-spacer_7 | pCRISPR-BEST with sgRNA:spacer7 | This study |
| pCRISPR-BEST-spacer_8 | pCRISPR-BEST with sgRNA:spacer8 | This study |
| pCRISPR-BEST-spacer_9 | pCRISPR-BEST with sgRNA:spacer9 | This study |
| pCRISPR-BEST-spacer_10 | pCRISPR-BEST with sgRNA:spacer10 | This study |
| pCRISPR-BEST-spacer_11 | pCRISPR-BEST with sgRNA:spacer11 | This study |
| pCRISPR-BEST-spacer_12 | pCRISPR-BEST with sgRNA:spacer12 | This study |

Supplementary Table 3. Primers used in this study.
*Forward primers were used as Sanger sequencing primers unless specifically indicated.*

| Names | sequence (5' to 3') | Purpose |
|---|---|---|
| ssDNA_spacer1 | CGGTTGGTAGGATCGACGGCAATGCCAGATTCTATTGATTGTTTTAGAGCTAGAAATAGA | For spacer cloning using ssDNA oligo bridging<br>For CRISPR-BEST validation |
| ssDNA_spacer2 | CGGTTGGTAGGATCGACGGCGCTCGGGATGATCATTTTGAGTTTTAGAGCTAGAAATAGA | |
| ssDNA_spacer3 | CGGTTGGTAGGATCGACGGCGCAGATGAGATTCAACTTATGTTTTAGAGCTAGAAATAGA | |
| ssDNA_spacer4 | CGGTTGGTAGGATCGACGGCGCTTCCGAATCAATAGAATCGTTTTAGAGCTAGAAATAGA | |
| ssDNA_spacer5 | CGGTTGGTAGGATCGACGGCCACATTGAAATCTGTTGAGTGTTTTAGAGCTAGAAATAGA | |
| ssDNA_spacer6 | CGGTTGGTAGGATCGACGGCTGGGCACATACCCTTTATCCGTTTTAGAGCTAGAAATAGA | |
| ssDNA_spacer7 | CGGTTGGTAGGATCGACGGCAATTCCGCTTAAATCCTCGAGTTTTAGAGCTAGAAATAGA | |
| ssDNA_spacer8 | CGGTTGGTAGGATCGACGGCCGAACCGGCACGAAAACTTGGTTTTAGAGCTAGAAATAGA | |
| ssDNA_spacer9 | CGGTTGGTAGGATCGACGGCACCACCGTTCTACAATGGAAGTTTTAGAGCTAGAAATAGA | |
| ssDNA_spacer10 | CGGTTGGTAGGATCGACGGCCCCGTTTTCATGGGGTTAATGTTTTAGAGCTAGAAATAGA | |
| ssDNA_spacer11 | CGGTTGGTAGGATCGACGGCGCTCGGGATGATCATTTTGAGTTTTAGAGCTAGAAATAGA | |
| ssDNA_spacer12 | CGGTTGGTAGGATCGACGGCGAACACGGCTTTGCACAAAGGTTTTAGAGCTAGAAATAGA | |
| Check_F1 | CTCGACACCATGATCGTGCG | For amplifying sequencing fragment containing spacers 1, 3, 4, 5, 7 |
| Check_R1 | CTCGTCGATCAGGGCGAAGT | |
| Check_F2 | TGGCTCGACCAGGACGTA | For amplifying sequencing fragment containing spacers 6, 10 |
| Check_R2 | AGACCTCCACCAGCAGGA | |
| Check_F3 | ATCGTATGGCATGAACGGGC | For amplifying sequencing fragment containing spacer 8, 11 |
| Check_R3 | ACGGTGATCCACTGGATG | |
| Check_F4 | TCCTTCAGTCGTTTCGGC | For amplifying sequencing fragment containing spacers 2, 9 |
| Check_R4 | GCAGCGACGTGTCGAACT | |
| Check_F5 | TACCACCTGCCCGTCAGGTA | For amplifying sequencing fragment containing spacers 12 |
| Check_R5 | GACCGTGCTGTCGTTCATCG | |
| Matrix_oligo_1 | CGGTTGGTAGGATCGACGGCGCTC1GC3AC5C6CAAGGACGCAGGTTTTAGAGCTAGAAATAGA | For spacer cloning using ssDNA oligo bridging |
| Matrix_oligo_2 | CGGTTGGTAGGATCGACGGCCGTTC2GC4AC6C7GGGGGACGCGGTTTTAGAGCTAGAAATAGA | |

| | | |
|---|---|---|
| Matrix_oligo_3 | CGGTTGGTAGGATCGACGGCAGCC1TC3GC5AC7TGCTCCAGTTGTTTTAGAGCTAGAAATAGA | For CRISPR-BEST characterization |
| Matrix_oligo_4 | CGGTTGGTAGGATCGACGGCCAAC1C2TC4GC6TCCCTCCTCCAGTTTTAGAGCTAGAAATAGA | |
| Matrix_oligo_5 | CGGTTGGTAGGATCGACGGCGATAC2C3TC5GC7CACATCGCATGTTTTAGAGCTAGAAATAGA | |
| Matrix_oligo_6 | CGGTTGGTAGGATCGACGGCCCGC1AC3C4TC6ACGGTTGGGTGGTTTTAGAGCTAGAAATAGA | |
| Matrix_oligo_7 | CGGTTGGTAGGATCGACGGCTCTGC2AC4C5TC7ACCCAACCGGGTTTTAGAGCTAGAAATAGA | |
| Matrix_test_1F | TTCGTGCGGAGCCGTTCATC | For amplifying sequencing fragment containing spacers of the Matrix |
| Matrix_test_1R | ACACGGGACTCGGTCACAGA | |
| Matrix_test_2F | ATCGACGACGCGGACTTCTC | |
| Matrix_test_2R | ATAGGCCGTTGAGGCGCTGA | |
| Matrix_test_3F | TCTACGAACTCACCTGGCCC | |
| Matrix_test_3R | GAACAGGCCCAGCAGGAGT | |
| Matrix_test_4F | GTACCTCGTCGGCACCATC | |
| Matrix_test_4R | CCGGTGAGCAGTTCCTCGT | |
| Matrix_test_5F | TATTCAGGGCGTACAGGTAG | |
| Matrix_test_5R | TGGGCGAACGTCGTCGAATT | |
| Matrix_test_6F | GATGGTCTCGACGGGACTC | |
| Matrix_test_6R | GGCATTCTGCTGACTCCGC | |
| Matrix_test_7F | AGTGTCAACGCGTGGCACG | |
| Matrix_test_7R | CTGTGCGCGTGCACCAGAT | |
| ssDNA_SCO5087-1 | CGGTTGGTAGGATCGACGGCCCGTTCACAGGTCGCGGCGGGTTTTAGAGCTAGAAATAGA | For spacer cloning using ssDNA oligo bridging<br>Spacers were from SCO5087 and SCO5092 |
| ssDNA_SCO5087-2 | CGGTTGGTAGGATCGACGGCCTACCGTTCACAGGTCGCGGGTTTTAGAGCTAGAAATAGA | |
| ssDNA_SCO5087-3 | CGGTTGGTAGGATCGACGGCGCCCTACCGTTCACAGGTCGGTTTTAGAGCTAGAAATAGA | |
| ssDNA_SCO5092 | CGGTTGGTAGGATCGACGGCGGTCCAGTCCGTGCACGTCGGTTTTAGAGCTAGAAATAGA | |
| 5087-check_F | CCGACGATGACGACGACCAC | For amplifying sequencing fragment containing spacers of SCO5087 or SCO5092 |
| 5087-check_R | CGCTGGGCACCAGGTAGTC | |
| 5092-check_F | GTTCGTTCGTGTCCGTCTCG | |
| 5092-check_R | GGCTGCTCAACCACCTGACC | |

| | | |
|---|---|---|
| ssDNA_kirN1 | CGGTTGGTAGGATCGACGGCGGGTTCCACGCGAACACGCCGTTTTAGAGCTAGAAATAGA | For spacer cloning using ssDNA oligo bridging<br>Spacer was from KirN |
| KirN_checkF | CGGCACGACCTCCCCTAC | For amplifying sequencing fragment containing KirN spacer |
| KirN_checkR | CGAACCGTTTCCATTCCGC | |
| stre_spacer_seq | TGTACGCGGTCGATCTTGA | For validation of spacer cloning |
| removalG_F | ACGGCCATGGTTTTAGAGC | For one G removal of sgRNA cassette from pCRISPR-Cas9 |
| removalG_R | GGAACATCGTAGCTGACG | |
| pHR1 | CGTGCGAUGCTAGCAAAGCGGTCGAT | For pCRISPR-Δ*kirN* construction and validation |
| pHR2 | AGGTGACCUCAGAACTCCATCTGGATTTGTTC | |
| pHR3 | ACAGGCGGUCGATCTTGACGGCTGGCG | |
| pHR4 | AGGTCTTCGUCGGCCGTCGATCCTACCAAC | |
| pHR5 | AGGTCACCUGAGGCGACCCCGACCAG | |
| pHR6 | ACCGCCTGUGCTCCTCCGTGTACGCGA | |
| pHR7 | ACGAAGACCUCGTCACGGTCCACTCCGA | |
| pHR8 | CACGCGAUATCTCCGACGACGCGTGG | |
| pHR9 | GGCGGCAACGCAGCG | |
| pHR10 | CGTACCGCTTCGGGCC | |
| pHR11 | GGACCGCCATGAGACCG | |
| pHR12 | GCGTGGTGGCCGGAG | |
| pHR13 | GTGCTCCTCCGTGTACGCG | |

Supplementary Table 4. The spacers used in this study.

| Names | Protospacer sequence (5' to 3') | PAM |
|---|---|---|
| Spacer1 | AATGCCAGATTCTATTGATT | CGG |
| Spacer2 | GCTCGGGATGATCATTTTGA | GGG |
| Spacer3 | GCAGATGAGATTCAACTTAT | TGG |
| Spacer4 | GCTTCCGAATCAATAGAATC | TGG |
| Spacer5 | CACATTGAAATCTGTTGAGT | AGG |
| Spacer6 | TGGGCACATACCCTTTATCC | GGG |
| Spacer7 | AATTCCGCTTAAATCCTCGA | AGG |
| Spacer8 | CGAACCGGCACGAAAACTTG | CGG |
| Spacer9 | ACCACCGTTCTACAATGGAA | CGG |
| Spacer10 | CCCGTTTTCATGGGGTTAAT | GGG |
| Spacer11 | GCTCGGGATGATCATTTTGA | AGG |
| Spacer12 | GAACACGGCTTTGCACAAAG | AGG |
| SCO5087-1 | CCGTTCACAGGTCGCGGCGG | AGG |
| SCO5087-2 | CTACCGTTCACAGGTCGCGG | CGG |
| SCO5087-3 | GCCCTACCGTTCACAGGTCG | CGG |
| SCO5092 | GGTCCAGTCCGTGCACGTCG | AGG |
| matrix_1 | GCTCGCACCCAAGGACGCAG | CGG |
| matrix_2 | CGTTCGCACCGGGGGACGCG | CGG |
| matrix_3 | AGCCTCGCACTGCTCCAGTT | CGG |
| matrix_4 | CAACCTCGCTCCCTCCTCCA | TGG |
| matrix_5 | GATACCTCGCCACATCGCAT | CGG |
| matrix_6 | CCGCACCTCACGGTTGGGTG | CGG |
| matrix_7 | TCTGCACCTCACCCAACCGG | TGG |
| KirN | GGGTTCCACGCGAACACGCC | GGG |

Table S5. sgRNA-kirN gBLOCK sequence.

| | |
|---|---|
| **sgRNA-*kirN* gBLOCK ® gene fragment** | GCTAGCAAAGCGGTCGATCTTGACGGCTGGCGAGAGGTGCGGGG AGGATCTGACCGACGCGGTCCACACGTGGCACCGCGATGCTGTTG TGGGCACAATCGTGCCGGTTGGTAGGATCGACGGACTAGTGATCG CATTTCGCCAACTACGTTTTAGAGCTAGAAATAGCAAGTTAAAATAA GGCTAGTCCGTTATCAACTTGAAAAAGTGGCACCGAGTCGGTGCTT TTTTTACCCTAGGAAAAGCTACGATGTTCCGGGGACTGCTGATCCG GTCAGCAGGTGGAAGAGGGACTGGATTCCAAAGTTCTCAATGCTG CTTGCTGTTCTTGAATGGGGGGTCGTTGACGACGACATGGCTCGAT TGGCGCGACAAGTTGCTGCGATTCTCACCAATAAAAAACGCCCGG CGGCAACGCAGCGTTCTGAACAAATCCAGATGGAGTTCTGAGGTC ACCTGAGG |

Supplementary Table 6. Detailed amino acid substitution list by CRISPR-BEST.
*Coding strand was masked with light blue, while non-coding strand was masked with light yellow.*

| AA | Codon | | AA |
|---|---|---|---|
| | *From* | *To* | |
| Ala | GCA | GTA | Val |
| | GCC | GCT | Ala |
| | GCC | GTC | Val |
| | GCC | GTT | Val |
| | GCG | GTG | Val |
| | GCT | GTT | Val |
| | GCA | ACA | Thr |
| | GCC | ACC | Thr |
| | GCG | ACA | Thr |
| | GCG | ACG | Thr |
| | GCG | GCA | Ala |
| | GCT | ACT | Thr |
| Arg | CGA | TGA | STOP |
| | CGC | TGT | Cys |
| | CGC | CGT | Arg |
| | CGC | TGC | Cys |
| | CGG | TGG | Trp |
| | CGT | TGT | Cys |
| | AGA | AAA | Lys |
| | AGG | AGA | Arg |
| | AGG | AAG | Lys |

| | | | |
|---|---|---|---|
| | AGG | AAA | Lys |
| | CGA | CAA | Gln |
| | CGC | CAC | His |
| | CGG | CAG | Gln |
| | CGG | CGA | Arg |
| | CGG | CAA | Gln |
| | CGT | CAT | His |
| Asn | AAC | AAT | Asn |
| Asp | GAC | GAT | Asp |
| | GAC | AAC | Asn |
| | GAT | AAT | Asn |
| Cys | TGC | TGT | Cys |
| | TGT | TAT | Tyr |
| | TGC | TAC | Tyr |
| Gln | CAA | TAA | STOP |
| | CAG | TAG | STOP |
| | CAG | CAA | Gln |
| Glu | GAG | AAA | Lys |
| | GAG | GAA | Glu |
| | GAG | AAG | Lys |
| | GAA | AAA | Lys |
| Gly | GGC | GGT | Gly |
| | GGA | AAA | Lys |
| | GGA | GAA | Glu |
| | GGA | AGA | Arg |
| | GGC | AAC | Asn |
| | GGC | AGC | Ser |
| | GGC | GAC | Asp |
| | GGG | AAA | Lys |
| | GGG | GAA | Glu |
| | GGG | AGA | Arg |
| | GGG | AAG | Lys |
| | GGG | GGA | Gly |
| | GGG | GAG | Glu |
| | GGG | AGG | Arg |
| | GGT | AAT | Asn |
| | GGT | AGT | Ser |
| | GGT | GAT | Asp |

| | | | |
|---|---|---|---|
| His | CAC | TAC | Tyr |
| | CAC | TAT | Tyr |
| | CAC | CAT | His |
| | CAT | TAT | Try |
| Ile | ATC | ATT | Ile |
| Leu | CTA | TTA | Leu |
| | CTC | TTT | Phe |
| | CTC | CTT | Leu |
| | CTC | TTC | Phe |
| | CTG | TTG | Leu |
| | CTT | TTT | Phe |
| | CTG | CTA | Leu |
| | TTG | TTA | Leu |
| Lys | AAG | AAA | Lys |
| Met | ATG | ATA | Ile |
| Phe | TTC | TTT | Phe |
| Pro | CCA | TTA | Leu |
| | CCA | TCA | Ser |
| | CCA | CTA | Leu |
| | CCC | TTC | Phe |
| | CCC | TCC | Ser |
| | CCC | TCT | Ser |
| | CCC | TTT | Phe |
| | CCC | CTT | Leu |
| | CCC | CCT | Pro |
| | CCC | CTC | Leu |
| | CCG | TTG | Leu |
| | CCG | TCG | Ser |
| | CCG | CTG | Leu |
| | CCT | TTT | Phe |
| | CCT | CTT | Leu |
| | CCT | TCT | Ser |
| | CCG | CCA | Pro |
| Ser | AGC | AGT | Ser |
| | TCA | TTA | Leu |
| | TCC | TTT | Phe |
| | TCC | TCT | Ser |
| | TCC | TTC | Phe |

| | | | |
|---|---|---|---|
| | TCG | TTG | Leu |
| | TCT | TTT | Phe |
| | TCG | TCA | Ser |
| | AGC | AAC | Asn |
| | AGT | AAT | Asn |
| Thr | ACA | ATA | Ile |
| | ACC | ACT | Thr |
| | ACC | ATC | Ile |
| | ACC | ATT | Ile |
| | ACG | ATG | Met |
| | ACT | ATT | Ile |
| | ACG | ACA | Thr |
| Trp | TGG | TGA | STOP |
| | TGG | TAA | STOP |
| | TGG | TAG | STOP |
| Tyr | TAC | TAT | Tyr |
| Val | GTC | GTT | Val |
| | GTA | ATA | Ile |
| | GTC | ATC | Ile |
| | GTG | ATA | Ile |
| | GTG | GTA | Val |
| | GTG | ATG | Met |
| | GTT | ATT | Ile |
| STOP | TGA | TAA | STOP |
| | TAG | TAA | STOP |

Supplementary Table 7. Frequency of STOP codon introduction in the nonessential secondary metabolites biosynthesis genes of *S. coelicolor* A3(2).

| ***S. coelicolor* A3(2)** | | | | |
|---|---|---|---|---|
| **Cluster no.** | **Cluster size (bp)** | **STOP count** | **gene count** | **STOP/gene** |
| 1 | 53018 | 498 | 40 | 12.45 |
| 2 | 24764 | 286 | 24 | 11.92 |
| 3 | 23530 | 302 | 22 | 13.73 |
| 4 | 49828 | 541 | 36 | 15.03 |
| 5 | 8242 | 78 | 6 | 13.00 |
| 6 | 38823 | 409 | 40 | 10.23 |
| 7 | 10399 | 86 | 9 | 9.56 |
| 8 | 10570 | 95 | 13 | 7.31 |
| 9 | 10972 | 112 | 9 | 12.44 |
| 10 | 79080 | 741 | 36 | 20.58 |
| 11 | 70903 | 740 | 63 | 11.75 |
| 12 | 20562 | 221 | 20 | 11.05 |
| 13 | 72543 | 640 | 70 | 9.14 |
| 14 | 10278 | 106 | 7 | 15.14 |
| 15 | 45282 | 408 | 35 | 11.66 |
| 16 | 11317 | 103 | 10 | 10.30 |
| 17 | 19321 | 204 | 16 | 12.75 |
| 18 | 13208 | 139 | 11 | 12.64 |
| 19 | 70203 | 862 | 33 | 26.12 |
| 20 | 46636 | 463 | 32 | 14.47 |
| 21 | 18700 | 200 | 18 | 11.11 |
| 22 | 25810 | 250 | 24 | 10.42 |
| 23 | 48144 | 540 | 37 | 14.59 |
| 24 | 26454 | 329 | 25 | 13.16 |
| 25 | 40888 | 419 | 36 | 11.64 |
| 26 | 21128 | 186 | 22 | 8.45 |
| 27 | 73251 | 713 | 65 | 10.97 |
| | | sum | sum | average |
| | | 9671 | 759 | 12.74 |

Supplementary Table 8. Frequency of STOP codon introduction in the nonessential secondary metabolites biosynthesis genes of *S. collinus* Tü365.

| ***S. collinus* Tü365** | | | | |
|---|---|---|---|---|
| **Cluster no.** | **Cluster size (bp)** | **STOP count** | **gene count** | **STOP/gene** |
| 1 | 23069 | 187 | 16 | 11.69 |
| 2 | 59480 | 525 | 45 | 11.67 |
| 3 | 162077 | 1381 | 72 | 19.18 |
| 4 | 25263 | 252 | 22 | 11.45 |
| 5 | 43867 | 431 | 32 | 13.47 |
| 6 | 20659 | 229 | 20 | 11.45 |
| 7 | 55714 | 655 | 31 | 21.13 |
| 8 | 11926 | 91 | 7 | 13.00 |
| 9 | 43264 | 482 | 34 | 14.18 |
| 10 | 41041 | 409 | 40 | 10.23 |
| 11 | 52568 | 483 | 38 | 12.71 |
| 12 | 10405 | 79 | 9 | 8.78 |
| 13 | 69134 | 648 | 59 | 10.98 |
| 14 | 10411 | 69 | 12 | 5.75 |
| 15 | 11770 | 113 | 9 | 12.56 |
| 16 | 42498 | 441 | 41 | 10.76 |
| 17 | 56291 | 594 | 34 | 17.47 |
| 18 | 55031 | 564 | 54 | 10.44 |
| 19 | 58662 | 564 | 45 | 12.53 |
| 20 | 21014 | 234 | 18 | 13.00 |
| 21 | 12169 | 122 | 7 | 17.43 |
| 22 | 12145 | 121 | 7 | 17.29 |
| 23 | 52987 | 555 | 47 | 11.81 |
| 24 | 11419 | 83 | 10 | 8.30 |
| 25 | 22163 | 195 | 18 | 10.83 |
| 26 | 13158 | 149 | 12 | 12.42 |
| 27 | 21011 | 170 | 19 | 8.95 |
| 28 | 26758 | 283 | 23 | 12.30 |
| 29 | 41374 | 354 | 37 | 9.57 |
| 30 | 73684 | 704 | 29 | 24.28 |
| 31 | 162077 | 1381 | 72 | 19.18 |
| 32 | 59480 | 525 | 45 | 11.67 |
| 33 | 23069 | 187 | 16 | 11.69 |
| | | sum | sum | average |
| | | 13260 | 980 | 13.53 |

### Supplementary Figure 1:

| In 5496474 - 5567376: | 1 | 2 | 3 | 4 | 5 | 6 | 7 | 8 | 9 | 10 | | PAM |
|---|---|---|---|---|---|---|---|---|---|---|---|---|
| | Hypothetic editing window | | | | | | | | | | | |
| Protospacer_1 | A | A | T | G | C | C | A | G | A | T | T C T A T T G A T T | CGG |
| | | | | | 29.5±5.4 | 92.9±10.1 | | | | | | |
| Protospacer_2 | G | C | T | C | G | G | G | A | T | G | A T C A T T T T G A | AGG |
| | | 0.00 | | 100.00 | | | | | | | | |
| Protospacer_3 | G | C | A | G | A | T | G | A | G | A | T T C A A C T T A T | TGG |
| | | 0.00 | | | | | | | | | | |
| Protospacer_4 | G | C | T | T | C | C | G | A | A | T | C A A T A G A A T C | TGG |
| | | 0.00 | | | 100.00 | 100.00 | | | | | | |
| Protospacer_5 | C | A | C | A | T | T | G | A | A | A | T C T G T T G A G T | AGG |
| | 0.00 | | 0.00 | | | | | | | | | |
| Protospacer_6 | T | G | G | G | C | A | C | A | T | A | C C C T T T A T C C | GGG |
| | | | | | 0.00 | | 90.7±8.5 | | | | | |
| Protospacer_7 | A | A | T | T | C | C | G | C | T | T | A A A T C C T C G A | AGG |
| | | | | | 100.00 | 100.00 | | 0.00 | | | | |
| Protospacer_8 | A | C | C | A | C | C | G | T | T | C | T A C A A T G G A A | CGG |
| | | | | | 70.1±10.8 | 70.1±10.8 | | | | 35.4±11.6 | | |
| Protospacer_9 | C | G | A | A | C | C | G | G | C | A | C G A A A A C T T G | CGG |
| | | | | | 100.00 | 100.00 | | | 0.00 | | | |
| Protospacer_10 | C | C | C | G | T | T | T | T | C | A | T G G G G T T A A T | GGG |
| | 0.00 | 0.00 | 0.00 | | | | | | 100.00 | | | |
| Protospacer_11 | A | C | C | G | T | T | C | C | A | T | T G T A G A A C G G | TGG |
| | | 0.00 | 0.00 | | | | 100.00 | 100.00 | | | | |
| Protospacer_12 | G | A | A | C | A | C | G | G | C | T | T T G C A C A A A G | AGG |
| | | | | 100.00 | | 100.00 | | | 0.00 | | | |

0 50 100

: Target C in coding strand

: Target C in non-coding strand

All sequences were displayed as 5'→3'

0 50 100

Color key of editing efficiency (%)

Supplementary Fig. 1. CRISPR-BEST evaluation by targeting twelve protospacers from the genome loci 5496474 – 5567376 of *S. coelicolor* A3(2). The entire 20nt protospacer plus 3nt PAM is displayed in 5' to 3' direction. The cytidines in the 10nt hypothetic editing window are labelled in red. The targeted C in coding and non-coding strand are masked as light yellow and light green, respectively. The editing efficiencies are shown as the mean and standard deviation. Five to ten exconjugants from each of the two to three independent conjugations were subjected to Sanger sequencing.

**Supplementary Figure 2:**

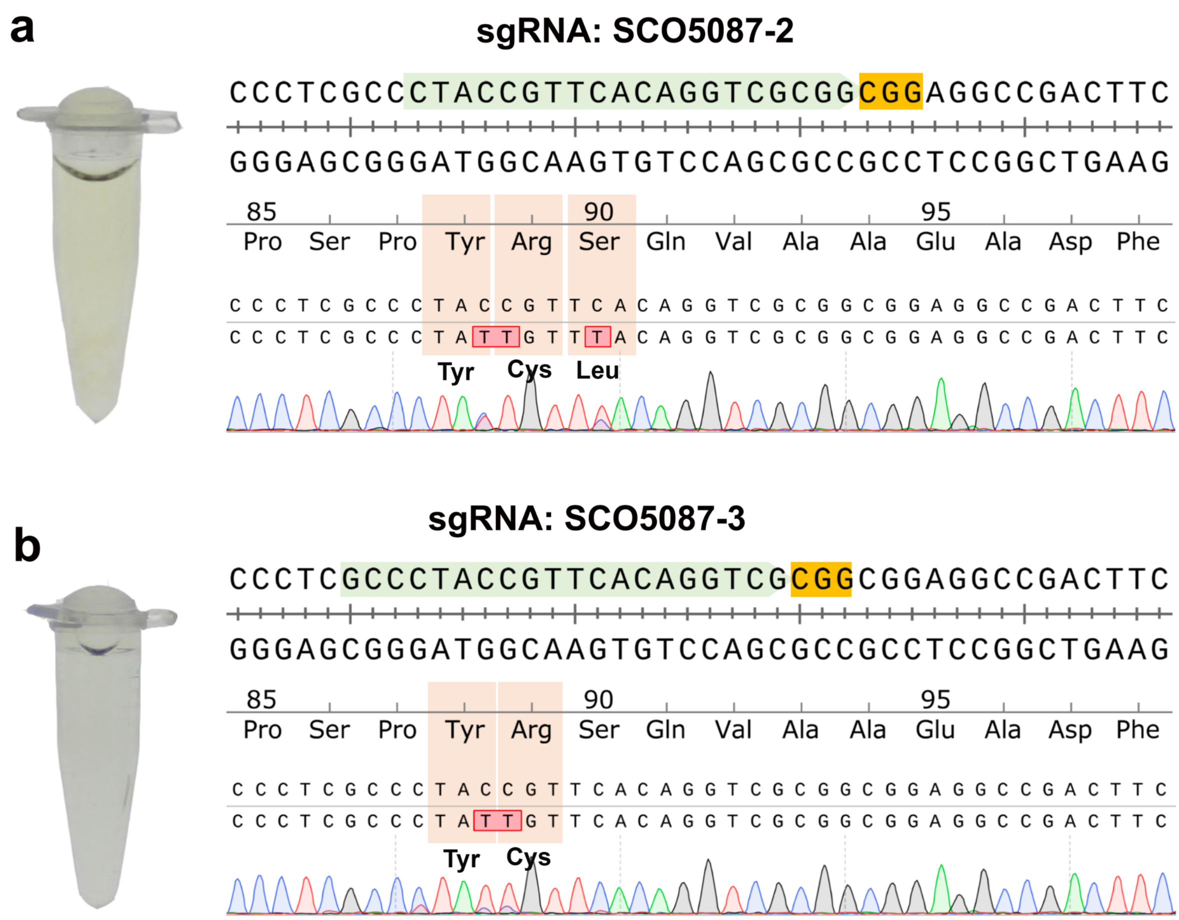

Supplementary Fig. 2. CRISPR-BEST editing validation by Sanger sequencing.

Sanger sequencing traces of the region containing a protospacer together with its PAM. Protospacers are highlighted in light green, PAM sequences in yellow, the codons and corresponding amino acids are indicated, detailed editing efficiency were shown in Supplementary Fig. 2a. **a.** the result of sgRNA SCO5087-2; **b.** the result of sgRNA SCO5087-3.

### Supplementary Figure 3:

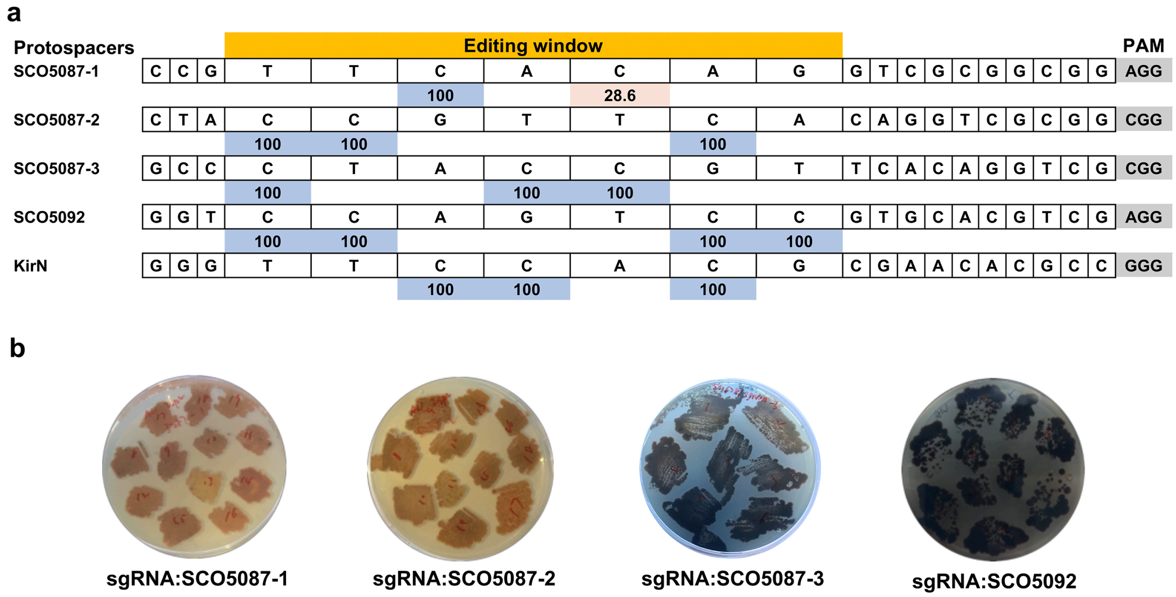

Supplementary Fig. 3. Editing efficiency of different protospacers.

**a.** The entire 20nt protospacer plus 3nt PAM is displayed. The 7nt editing window is shown in yellow. The numbers indicate the editing efficiency of the target cytidine. **b.** seven to twelve of the exconjugants with different sgRNAs were randomly picked and streaked onto the apramycin and nalidixic acid containing ISP2 plates, and incubated for five days at 30 °C. The photos were taken by a ColonyDoc-It™ Imaging Station (Analytik Jena AG, Germany) at day 6.

### Supplementary Figure 4:

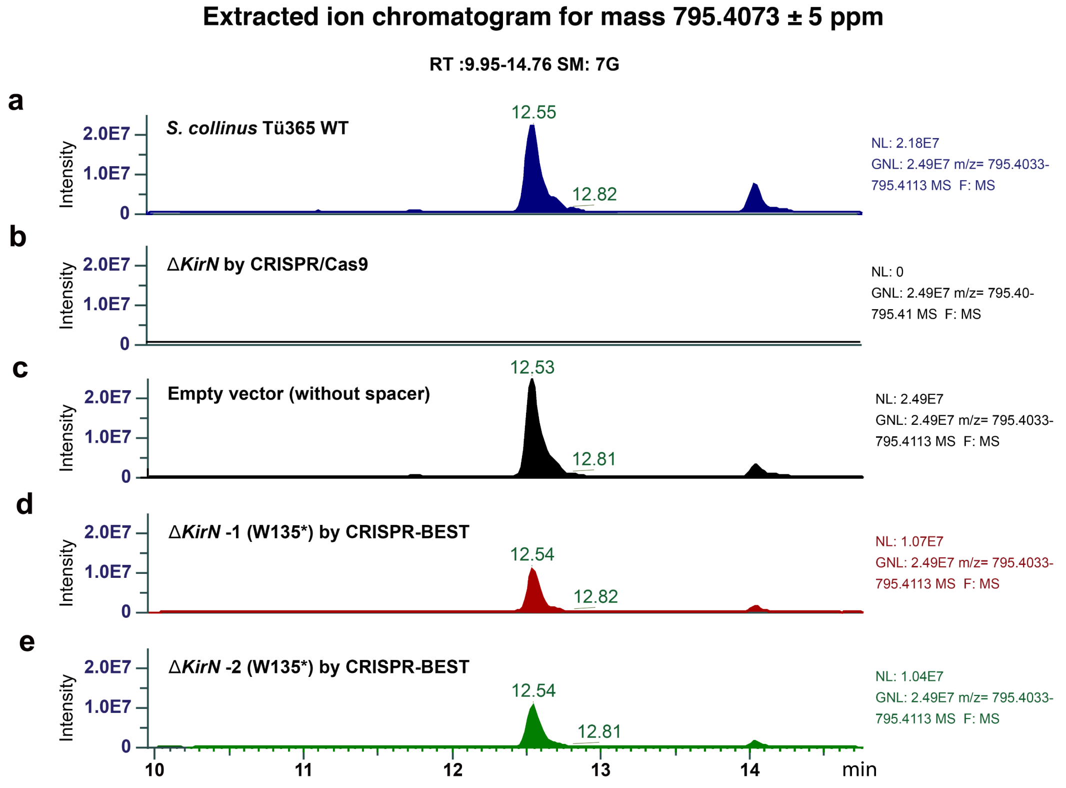

Supplementary Fig. 4. Chemical analysis of the extracts from shake flask fermentation, UV-Vis profiles of **a.** WT *S. collinus* Tü365, **b.** Δ*kirN*-Cas9 (+undesired deletion of chromosomal arms), **c.** *S. collinus* Tü365 with pCRISPR-BEST empty vector (without spacer), and **d.** and **e.** two *kirN*$_{W135\rightarrow STOP}$ mutants generated by CRISPR-BEST were shown.

**Supplementary Figure 5:**

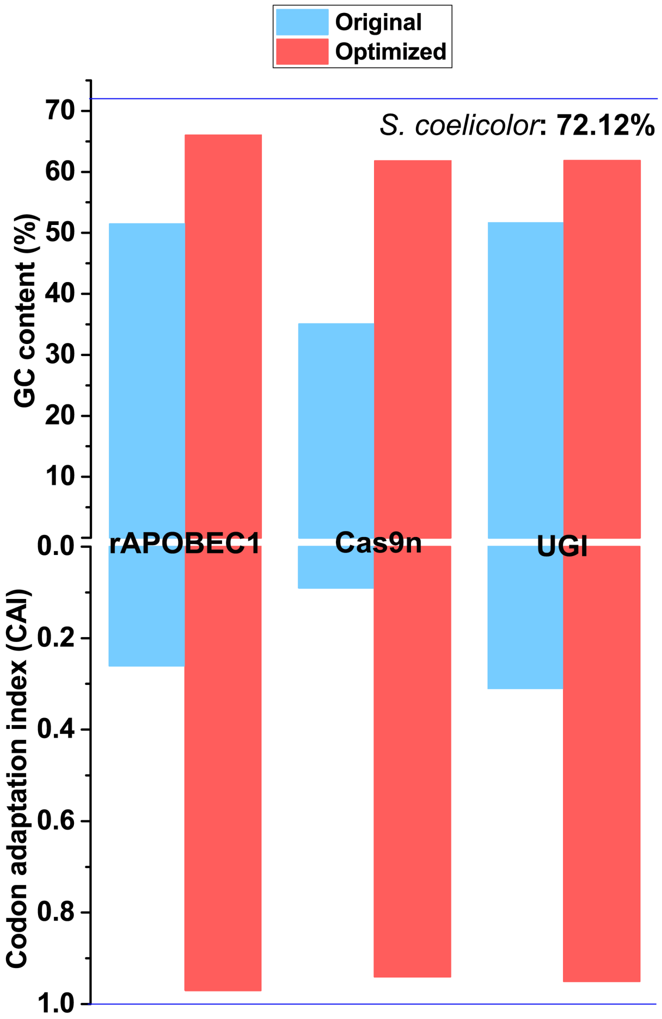

Supplementary Fig. 5. Codon optimization results of the three proteins.

**Supplementary Figure 6:**

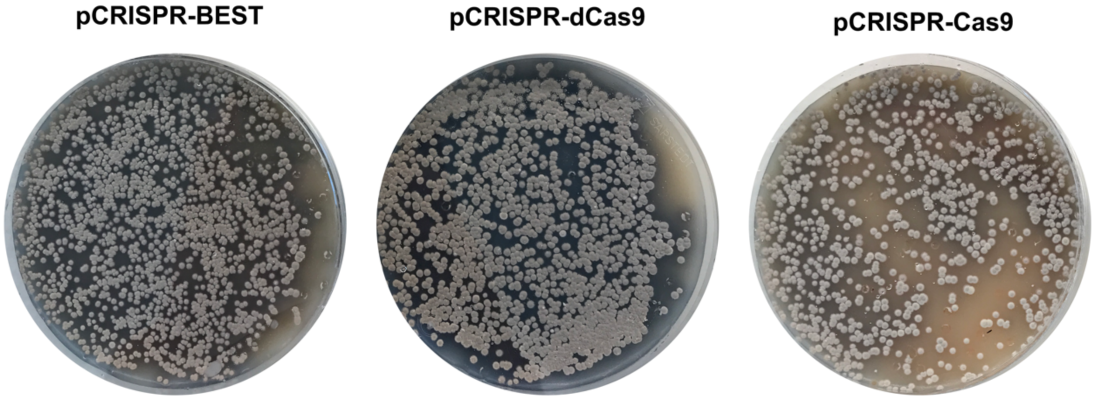

Supplementary Fig. 6. The conjugation results of three plasmids carrying sgRNA from the same region.

### Supplementary Figure 7:

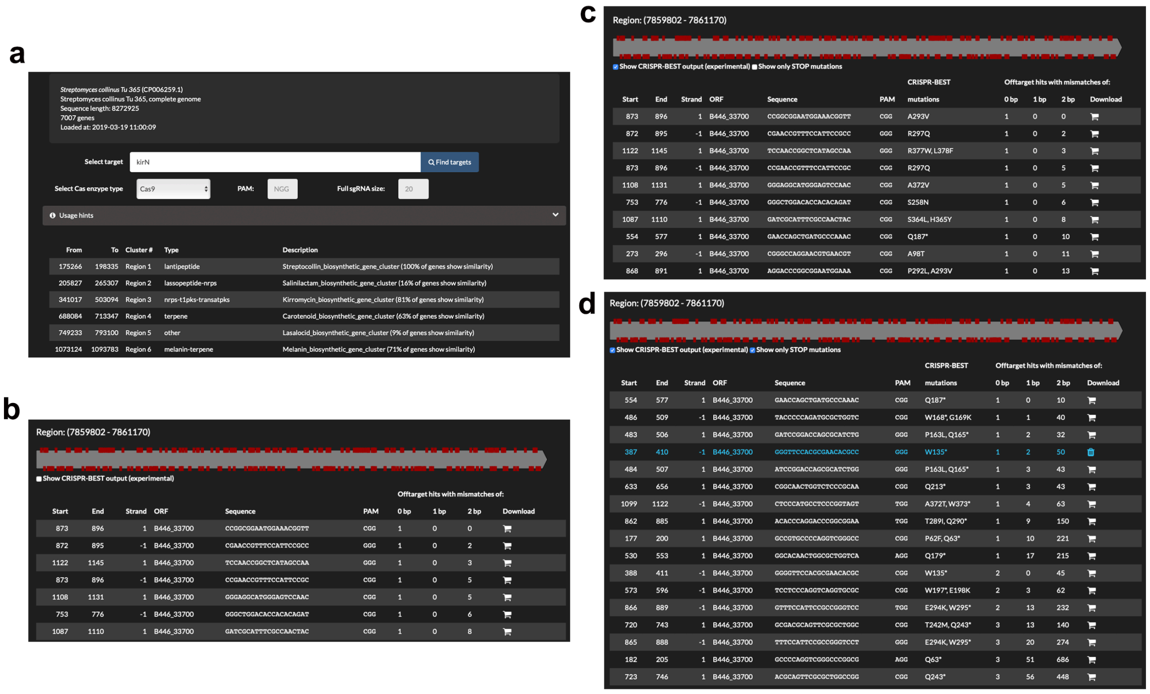

Supplementary Fig. 7. An example output of CRISPy-web run to identify CRISPR-BEST compatible protospacers in the *kirN* gene of *S. collinus* Tü365. **a.** Overview of protospacers identified in the BGCs of *S. collinus* Tü365. **b.** Zoom view of *kirN*. **c.** Selected view of CRISPR-BEST compatible protospacers. **d.** Selected view of protospacers that can introduce STOP codons into *kirN* gene.
